# Genetic differentiation in the *MAT*-proximal region is not sufficient for suppressing recombination in *Podospora anserina*

**DOI:** 10.1101/2024.12.03.626592

**Authors:** Pierre Grognet, Robert Debuchy, Tatiana Giraud

## Abstract

Recombination is advantageous over the long-term, as it allows efficient selection and purging deleterious mutations. Nevertheless, recombination suppression has repeatedly evolved in sex chromosomes and mating-type chromosomes. The evolutionary causes for recombination suppression and the proximal mechanisms preventing crossing overs are still poorly understood. Several hypotheses have recently been suggested based on theoretical models, and in particular, that divergence could accumulate neutrally around a sex-determining region and reduce recombination rates, a self-reinforcing process that could foster progressive extension of recombination suppression. The ascomycete fungus *Podospora anserina* is an excellent model for investigating these questions. A 0.8 Mb region around the mating-type locus is non-recombining, despite being collinear between the two mating types. This fungus is mostly selfing, so that strains are highly homozygous, except in the non-recombining region around the mating-type locus that displays differentiation between mating types. Here, we generated a mutant to test the hypothesis that sequence divergence alone is responsible for recombination cessation. We replaced the *mat-*idiomorph by the sequence of the *mat+* idiomorph, to obtain a strain that is sexually compatible with the *mat-*reference strain and isogenic to this strain in the *MAT*-proximal region. Crosses showed that recombination was still suppressed in the *MAT*-proximal region in the mutant strains, indicating that other proximal mechanisms than inversions or mere sequence divergence are responsible for recombination suppression in this fungus. This finding suggests that selective mechanisms likely acted for suppressing recombination, as the neutral model does not seem to hold, at least in this fungus.

**ARTICLE SUMMARY:** In many organisms, a non-recombining region is observed around the sex-determining locus. In the fungus *Podospora anserina*, recombination is suppressed within an 800kb region around the mating-type locus. In natural strains, the genome is isogenic between two mating-types, except in this region displaying heterozygosity. To determine if this heterozygosity can be responsible for the lack of recombination, compatible strains were engineered to be isogenic, including in the non-recombining region. Recombination inhibition persisted, indicating that mere sequence divergence does not cause recombination suppression. Our study provides an interesting insight on the molecular and evolutionary causes of recombination inhibition.

## INTRODUCTION

Recombination is widespread in eukaryotes, as it is advantageous over the long-term. Recombination breaks up allelic combinations, which allows more efficient selection and the purging of deleterious mutations (Muller 1932; Hill and Robertson 1966). Nevertheless, recombination can be suppressed locally in genomes, the most studied cases being on sex chromosomes (Ellegren 2000; Bachtrog 2014; Beukeboom and Perrin 2014; Cortez *et al*. 2014; Ma and Veltsos 2021). Recombination suppression has repeatedly evolved in sex chromosomes, although the reason why is still debated (Wright *et al*. 2016; Abbott *et al*. 2017; Ponnikas *et al*. 2018; Charlesworth 2021; Jay *et al*. 2024; Saunders and Muyle 2024). Indeed, while sexual antagonism has long been a commonly accepted hypothesis to explain progressive extension of recombination cessation on sex chromosomes, for linking to sex-determining genes other genes with alleles beneficial in only one of the sexes (Charlesworth *et al*. 2005; Charlesworth 2021), little evidence could be found in favor of this hypothesis (Ironside 2010). Furthermore, repeated evolution of recombination suppression has also been shown on fungal mating-type chromosomes, despite the lack of sexual antagonism (Fraser *et al*. 2004; Menkis *et al*. 2008; Branco *et al*. 2017; Branco *et al*. 2018; Bazzicalupo *et al*. 2019; Hartmann, Duhamel, *et al*. 2021; Hartmann, Ament-Velásquez, *et al*. 2021; Duhamel *et al*. 2022; Vittorelli *et al*. 2023; Jay *et al*. 2024). Indeed, there are no sex roles, or other obvious differentiated traits, associated to mating types in fungi and that would be controlled by genes distinct from the mating-type genes themselves (Branco *et al*. 2017; Bazzicalupo *et al*. 2019; Hartmann, Duhamel, *et al*. 2021).

Other hypotheses than sexual antagonism have therefore been proposed to explain the evolution of recombination suppression on sex-related chromosomes (Wright *et al*. 2016; Abbott *et al*. 2017; Kent *et al*. 2017; Ponnikas *et al*. 2018; Jeffries *et al*. 2021; Jay *et al*. 2022; Jay *et al*. 2024; Saunders and Muyle 2024). It has been suggested that there could be a selection of non-recombining fragments that would carry fewer deleterious mutations than average, and that the few recessive deleterious mutations would be sheltered at the heterozygous stage when associated to a Y-like sex-determining or mating-type determining gene, allowing the fixation of non-recombining fragments despite their load (Jay *et al*. 2022; Lenormand and Roze 2022; Jay *et al*. 2024). This would correspond to an evolutionary cause selecting against recombination, while the proximal mechanism preventing recombination could be inversions or any other mechanism, from *cis*- or *trans*-acting recombination modifiers to epigenetic marks. Another hypothesis involves recombination-suppressing epigenetic marks, possibly associated with transposable elements for their silencing, known to accumulate in non-recombining regions, and that could spread in nearby regions (Kent *et al*. 2017). A related hypothesis postulates that sequences accumulate differences between sex-related chromosomes at the margin of a sex- or mating-type locus due to linkage disequilibrium, and that such decrease in sequence similarity reduces recombination rates, which further decreases sequence identity as a self-reinforcing process (Jeffries *et al*. 2021). These later hypotheses correspond to both the evolutionary and proximal cause of recombination suppression.

The ascomycete fungus *Podospora anserina* is an excellent model for investigating the questions of the evolutionary and proximal causes of recombination suppression. Early genetic analyses showed a peculiar pattern of recombination on the mating-type bearing chromosome (chromosome 1) (Marcou *et al*. 1979), indicating second-division segregation of the mating-type (*MAT*) locus due to an obligate crossover between the *MAT* locus and the centromere of chromosome 1. Several markers were found to be tightly linked to the *MAT* locus, suggesting the existence of a non-recombining region. Later, a 0.8 Mb non-recombining region around the mating-type locus have been described (Grognet, Bidard, *et al*. 2014). There can however be some very rare events of recombination in this region (Contamine *et al*. 1996). The non-recombining region is collinear between the two mating types (Grognet, Bidard, *et al*. 2014), indicating that the proximal causes for the lack of crossing-overs are not inversions or other genomic rearrangements. A localized hotspot of concentrated repeats within the *MAT*-proximal region was suspected to play a role, but its deletion did not restore recombination (Grognet, Bidard, *et al*. 2014). Regarding the evolutionary cause of recombination suppression, sexual antagonism cannot apply as the mating-type chromosomes do not control any differences in gamete size or behavior (Silar 2020). Furthermore, the haploid phase, in which cells are of alternative mating types, is virtually nonexistent in *P. anserina*, as ca. 99% of sexual spore produced after meiosis are already carrying two nuclei of opposite mating types (Marcou *et al*. 1979), a feature that is faithfully maintained in the mycelium (Grognet, Bidard, *et al*. 2014). There is therefore little room in the life cycle for antagonistic selection between mating types.

*P. anserina* is mostly selfing, by automixis, so that strains are typically highly homozygous, except in the non-recombining region around the mating-type locus, that displays differentiation between mating types (98.44% identity) (Grognet, Bidard, *et al*., 2014; Hartmann *et al*., 2021). Here, we generated a mutant to test the “neutral hypothesis” postulating that divergence alone is responsible for recombination cessation in the *MAT*-proximal region (Jeffries *et al*. 2021). We replaced the *mat*-sequence at the *MAT* locus by the sequence of the *mat*+ idiomorph, to obtain a *mat+* strain isogenic to the *mat-*strain in the non-recombining *MAT*-proximal region. We crossed the *mat*+ mutant with the *mat*-wild-type strain, obtaining a dikaryotic strain homozygous in the *MAT*-proximal region. We used strains with resistance genes as markers to detect recombination events. We analyzed the progenies of several crosses, which showed that recombination was still suppressed in the *MAT*-proximal region in the strain homozygous in the *MAT*-proximal region. These findings indicate that other proximal mechanisms than divergence is responsible for recombination suppression in this fungus.

## MATERIAL AND METHODS

### Strains and media

The strain used for transformation is deleted for *Ku70* (El-Khoury *et al*. 2008). In that strain, the *ku70* gene has been replaced by a geneticin resistance cassette.

### Plasmids

The plasmid containing the *mat*+ locus sequence comes from the *P. anserina* 10 kbp plasmid genomic library (Espagne *et al*. 2008). Plasmid GAOAB40BC11 contains the *mat*+ locus (3842 bp) with 2.9 kbp upstream and 3.9 kbp downstream. The plasmid was linearized by *Nar*I before transformation. *Nar*I is absent from the *P. anserina* sequences cloned in GAOAB40BC11 and present in the vector sequences of this plasmid.

Since the transformation is made in a NHEJ defective strain, the plasmid containing the hygromycine resistance cassette used for co-transformation needs to integrate via homologous recombination. To do so, we cloned the *Pa_2_3690* gene sequence into the pBC-Hygro plasmid. The gene *Pa_2_3690* is genetically independent from the *MAT* locus, its inactivation does not lead to abnormal phenotype (Ait Benkhali *et al*. 2013) and it has already been used for targeted integration (Déquard-Chablat *et al*. 2012). The *Pa_2_3690* gene sequence (1935 bp) has been PCR amplified using primers pBCpro3690 (ggccgctctagaactagtggatcccccTCACTCCACAAGGCAGCATCTAA) and pBCter3690 (aagcttgatatcgaattcctgcagcccCAGGTGCAAAGGTAACACTCGGT). The purified PCR product has been cloned into pBC-Hyg linearized by *Sma*I using NEB’s NEBuilder HiFi DNA Assembly to give plasmid pBCH3690.

### Transformation

Three transformations were performed, yielding 18, 33 and 158 hygromycin resistant transformants, respectively. The mating-type phenotypes of the hygromycin-resistant transformants were tested by confronting each of them with tester strains of known mating types. From the first transformation, one transformant was able to mate only with the *mat*-tester strain, indicating that its mycelium contained only *mat*+ nuclei, and therefore that the replacement of the mating-type idiomorph was successful. Another transformant could mate with both tester strains, suggesting that it contained *mat*+ and *mat*-nuclei; a mixture of *mat+* and *mat-*nuclei is not surprising as *P. anserina*’s protoplasts used for transformation can contain several nuclei. The remaining 16 transformants mated only with the *mat+* tester strain, indicating transformation with only the hygromycin resistance cassette but not the mating-type idiomorph. The two other transformation attempts gave respectively zero and six *mat*+ transformants, three and 109 transformants with both mating types, and 30 and 39 *mat*-transformants. From the last transformation assay, four transformants were not able to mate with any of the tester strains. The following steps are described in the result section.

### Sequencing

Amplifications of sequences upstream and downstream of the *mat+* idiomorph of the *LPRM* strain were performed with the two primer pairs gggacctctgcagggaat/cactggaacggaggagga and tgacgaatgaaatcgtcgaa/gacccaccgaacctcctc. The genome of strain *LPRM* has been sequenced using Illumina technology (paired-end 150bp at Novogene). The reads have been mapped on both *mat+* and *mat-*genome sequences using bowtie2 (version 2.5.1). Mutations have been called using samtools mpileup (version 1.9) and processed with custom-made R scripts. Around the mating-type locus, the sequence has been confirmed by visual inspection of the mapped reads.

Surprisingly, three small possible deletions (170 bp, 55 bp and 240 bp long, respectively) were detected on chromosome 2 compared to the reference genome published for the *S* strain (Espagne *et al*. 2008). The two first ones were in the coding sequences of putative genes: Pa_2_1350, which has no predicted domain and is present only in *Podospora* species or close relatives, and Pa_2_3445, which has no predicted function but carries a kinesin domain and is conserved in ascomycete fungi. None of these two genes seem to be expressed in the available RNA-seq data (Lamacchia *et al*. 2016; Benocci *et al*. 2018; Silar *et al*. 2019; Lelandais *et al*. 2022). The third small deletion was in an intergenic region between the genes Pa_2_6390 and Pa_2_6380. To understand these deletions, we checked the aligned reads of previous sequencing data of our wild-type *S* strain (ChIP-seq data, (Carlier *et al*. 2021)). These three putative deleted sequences were also devoid of aligned reads in these samples (Figure S1), suggesting errors in the reference genome sequence or that these mutations occurred in our wild-type *S* strain prior to the experiments performed here. The rest of the genome do not show any other mutation, except a few C:G/T:A substitutions on repeated sequences that might be caused by RIP (repeat-induced point mutations (Gladyshev 2017)).

### Statistical analyses

The 2×2 contingency tests were performed with the GraphPad software (https://www.graphpad.com/quickcalcs/contingency1), with Fisher’s exact tests for small sample size and two tailed p-values.

## Results and Discussion

### Mating-type locus replacement

In order to get sexually compatible strains with isogenic *MAT*-proximal regions, we replaced the *MAT* locus of a *mat*-strain by the *mat*+ idiomorph. To do so, a linearized plasmid containing the *mat*+ idiomorph sequence has been co-transformed with a plasmid containing a hygromycin resistance cassette in a NHEJ defective strain (*i.e*. unable of non-homologous end joining) (El-Khoury *et al*. 2008), to allow only integration by homologous recombination at the mating-type locus. Integration of the *mat*+ containing plasmid thanks to two recombination events leads to the replacement of the *MAT* locus (Figure 1). The hygromycin resistance cassette allows the selection of the transformant as co-transformation is efficient in *P. anserina*: a plasmid containing a resistance gene is mixed with the replacement cassette (five to ten time more of the cassette) prior transformation. In such conditions, most hygromycin-resistant transformants will here be expected to have also integrated the cassette at the *MAT* locus, which can then be checked. The plasmid containing the hygromycine resistance marker was targeted to the gene Pa_2_3690, which was previously identified as an optimal target for genomic integration of any DNA fragment (Déquard-Chablat *et al*. 2012). Three transformations were performed, yielding 209 hygromycin resistant transformants in total. The mating-type phenotypes of the hygromycin-resistant transformants were tested by confronting each of them with tester strains of known mating types. From two independent transformations, seven transformants were able to mate only with the *mat*-tester strain, indicating that their mycelium contained only *mat*+ nuclei, and therefore that the replacement of the mating-type idiomorph was successful.

**Figure 1:**
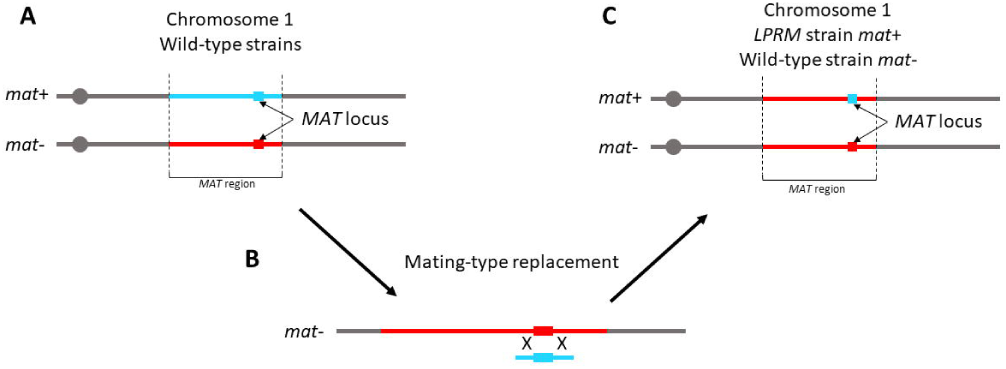
Strategy for the mating-type locus replacement: A) The right arm of chromosome 1 for both *mat*+ and *mat*-wild-type strain is depicted (not to scale). The *MAT* region is the 0.8 Mpb region devoid of recombination. The squares represent the *MAT* loci containing the mating-type genes. B) The *mat-*strain has been transformed with a plasmid containing the mat+ sequence and a plasmid containing the hygromycin resistance marker. This latter plasmid was targeted to a gene previously identified as an optimal target for genomic integration of any DNA fragment (https://doi.org/10.4148/1941-4765.1011). Thanks to two recombination events, the *MAT* was replaced, generating the *LPRM* strain. C) The *LPRM* strain (Locus Plus - Region Minus) can be crossed with a wild-type *mat-*strain. In this context, the *MAT* regions are strictly identical.

Five of the *mat*+ transformants from the different transformation assays were crossed with the *mat*-wild-type strain, and homokaryotic spores from the progenies have been isolated. Subsequent crossing of these progenies allowed us to recover a NHEJ proficient strain without the hygromycin-resistance cassette harbored by the plasmid used for co-transformation. Hence, the *mat*+ individuals from these progenies carry the *mat*+ idiomorph sequence but the rest of the genome should be identical to the *mat*-strain, including in the *MAT*-proximal region. One of these strains, from now on called *LPRM* (Locus Plus - Region Minus, Figure 1), was randomly selected and used for further investigation.

### Sequence analyses of the mutant strain

In order to have a first confirmation of the replacement of the *MAT* locus and to localize the recombination events, we first PCR amplified the regions directly adjacent to the mating-type locus on both sides and sequenced the 800 bp PCR amplicons. We searched within these sequences for single nucleotide polymorphisms (SNPs) and indels previously identified between the wild-type *mat*+ and *mat*-strains (Grognet, Bidard, *et al*. 2014). On one side (toward the centromere), the sequence obtained was identical to the sequence of the *mat*+ strain up to 80 bp away from the *MAT* locus, but had a *mat-*allele at the next SNP, 247 bp away from the *MAT* locus (Fig. 2), indicating that the recombination with the cassette occurred between these two positions. On the other side, only one SNP was present and it displayed the *mat*+ allele. To further check the locus replacement, we sequenced the whole genome of the *LPRM* strain. We looked at the sequence around the *MAT* locus and focused on SNPs and indels (Figure 2). The genome sequence confirmed that the recombination events replacing the *MAT* locus took place as expected, on one side between position -80 and -247 (in bp, relative to the border of the *MAT* locus), as the site at -80 shows a *mat*+ genotype and the site at -247 shows a *mat*-genotype. On the other side, sites at position +31 and +1334 both carried a *mat+* genotype, whereas sites at +5419 and further away were all of *mat-*genotype. The allele of the site at position +1617 could not be accurately determined due a long stretch of C and a poor read coverage at that position. The rest of the *MAT*-proximal region carried the *mat*-specific base pairs, indicating that the replacement of the *mat-*idiomorph occurred without affecting the chromosomal structure of the *MAT*-proximal region.

**Figure 2:**
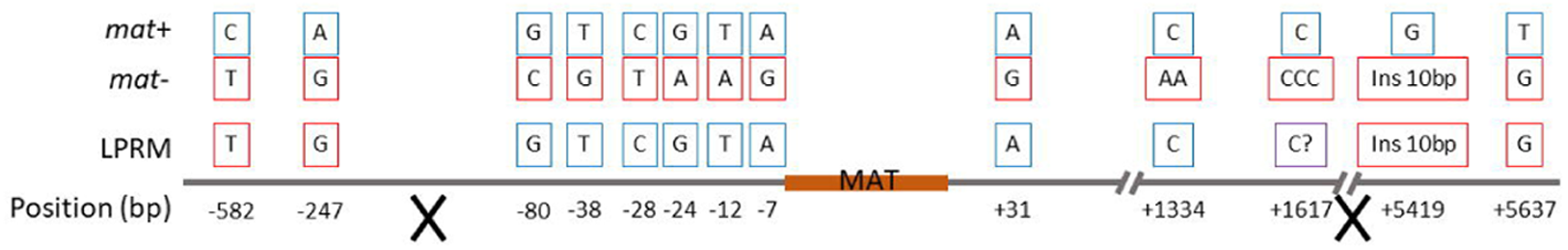
Sequence polymorphism analysis around the *MAT* locus. Base pairs specific to *mat*+ (blue) and *mat*-(red) are shown on top and their position relative to the *mat*+ idiomorph at the bottom. The sequence of the *LPRM* strain carries *mat+* SNPs until 80 bp downstream of the *mat+* idiomorph and *mat-*SNPs further away. On the other side, *mat*+ SNPs are found until 1617 bp upstream of the *mat+* idiomorph and *mat-*SNPs and indels are present from the next polymorphism at 5419 bp. The crosses indicate the localization of the recombination events. Ins= insertion.

### The mutant *LRPM* strain phenotype is similar to the wild type

The *LPRM* strain phenotype was indistinguishable from the one of the wild-type *S* strain. When grown as monokaryon, the mycelium had the same growth rate, pigmentation and overall aspect as the *S* strain (data not shown). We also compared the phenotype of a heterokaryon (which is the dominant life stage of *P. anserina*) of the *S* strain (*mat*+ and *mat*-) with a heterokaryon resulting from the vegetative fusion of *LPRM* and *S mat*-(Figure 3). The two heterokaryons were again indistinguishable: they both formed the typical ring of perithecia, the perithecia were equally numerous, properly shaped and the spores were normally formed and in similar amounts. These observations suggest that, at least in our laboratory conditions, the homozygosity at the *MAT*-proximal region has no obvious effects on vegetative or reproductive traits.

**Figure 3:**
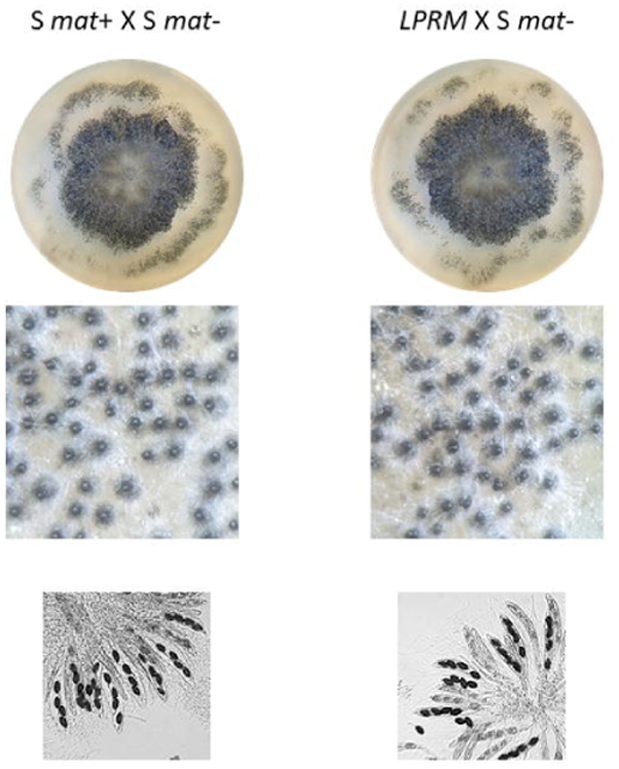
*LPRM* strain phenotype. The *LPRM* strain was compared to the *S mat*+ strain in heterokaryon with the *S mat*-strain. These two heterokaryons were indistinguishable in terms of mycelium growth (upper pictures), perithecium formation (middle pictures) or spore formation (lower pictures).

### Crosses indicate suppressed recombination in the mutant strain homozygous in the *MAT*-Proximal region

To test whether the inhibition of recombination still occurs in a cross of *LPRM* with *S mat*-despite the homozygosity in the *MAT*-proximal region, we took advantage of two strains already available with hygromycin resistance markers in the *MAT* region, to detect recombination events (Figure 4A). The Δ*PaRid* strain (Grognet *et al*. 2019) had its CDS (coding sequence) of the *PaRid* gene replaced by a hygromycin resistance cassette. The Δ*Pa_1_18960* strain (Grognet, Bidard, *et al*. 2014) had its CDS of the *Pa_1_18960* gene replaced by a hygromycin resistance cassette. These two strains can be crossed and previous work showed that there is no recombination between these loci and the *MAT* locus. Hence, when crossing the two mutant strains [Hygro^R^, mat-] with *S mat*+ and *LPRM*, recombination events in the *MAT*-proximal region should produce [Hygro^R^, mat+] and [Hygro^S^, mat-] progeny and can thereby be easily detected. We made these four crosses twice (each of the two mutant strains [Hygro^R^, mat-] with *S mat*+ and *LPRM*), and collected homokaryotic spores from each cross. We looked for recombinants in the homokaryotic progeny by determining i) the mating type with tester strains and ii) hygromycin resistance on selective medium. The results are shown in Table 1. When *S mat*+ was crossed with either Δ*PaRid* or Δ*Pa_1_18960,* no recombinants in the *MAT*-proximal region were recovered across 174 and 188 analyzed offspring, respectively. When *LPRM* was crossed with Δ*PaRid,* no recombinants were recovered from 210 offspring, and when crossed with Δ*Pa_1_18960,* a single recombinant was found out of 222 offspring (0.45 %). Given the distance between the *MAT* locus and the two other loci, we could expect much more recombination events if the recombination were fully restored (see below).

**Figure 4:**
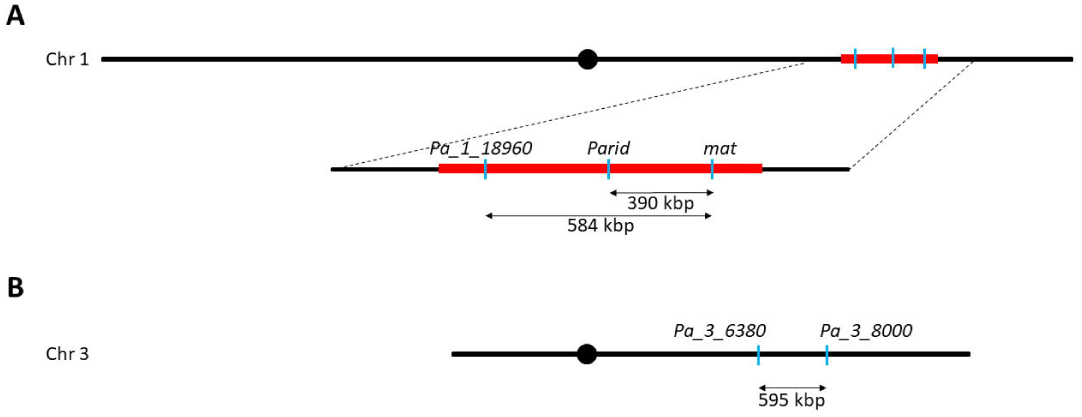
Schematic representation of A) chromosome 1 and B) chromosome 3. Chromosome sizes are scaled as well as the position of the centromere (black disk), the loci of the markers used in the study (blue line), and position and size of the non-recombining *MAT*-region (in red). Physical distances are given below the double headed arrows.

**Table 1:**
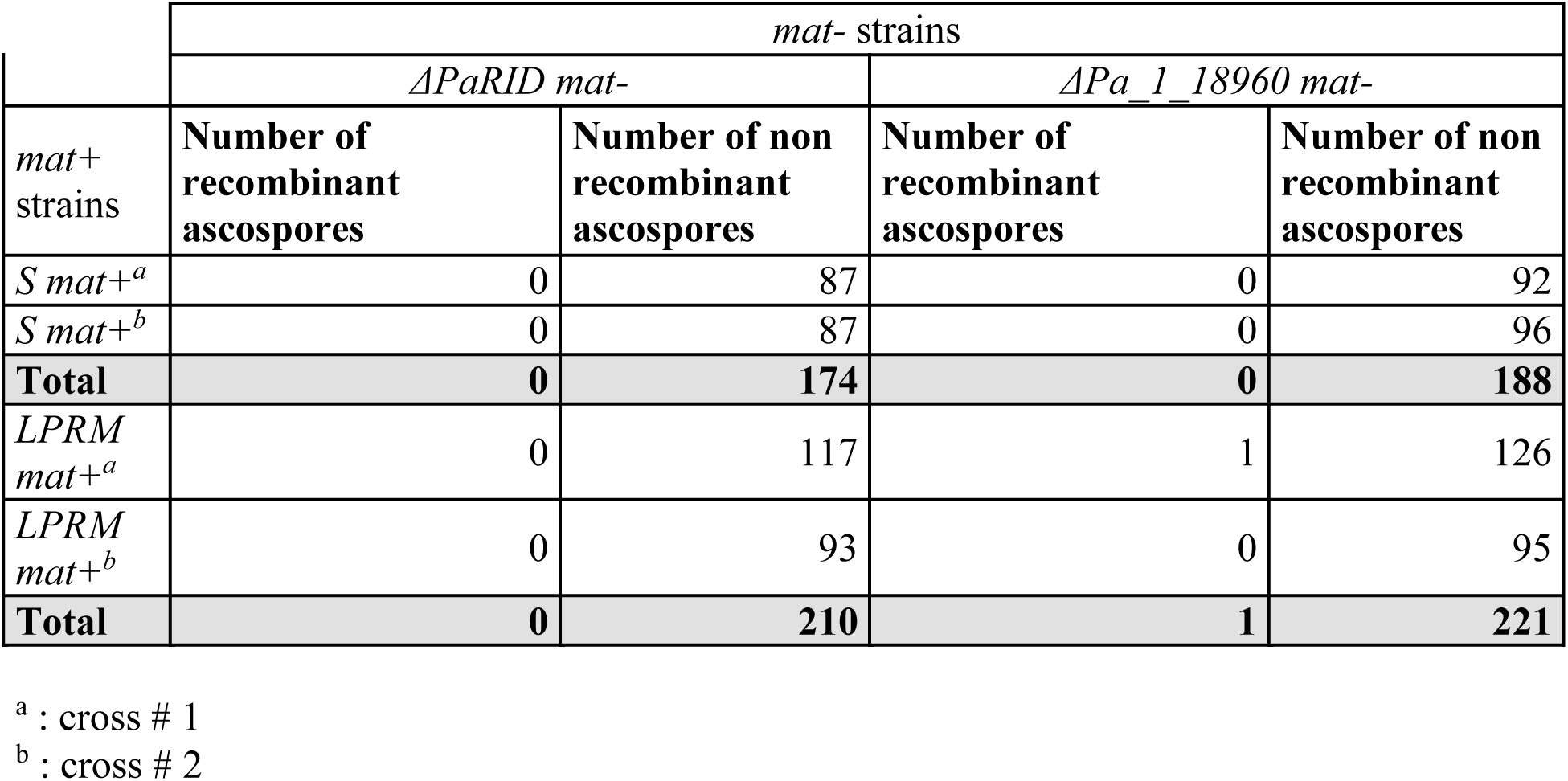
Number of offspring showing recombination or no recombination between markers near the mating-type locus and the mating-type locus, in wild type and LPRM contexts.

**Table 2:**
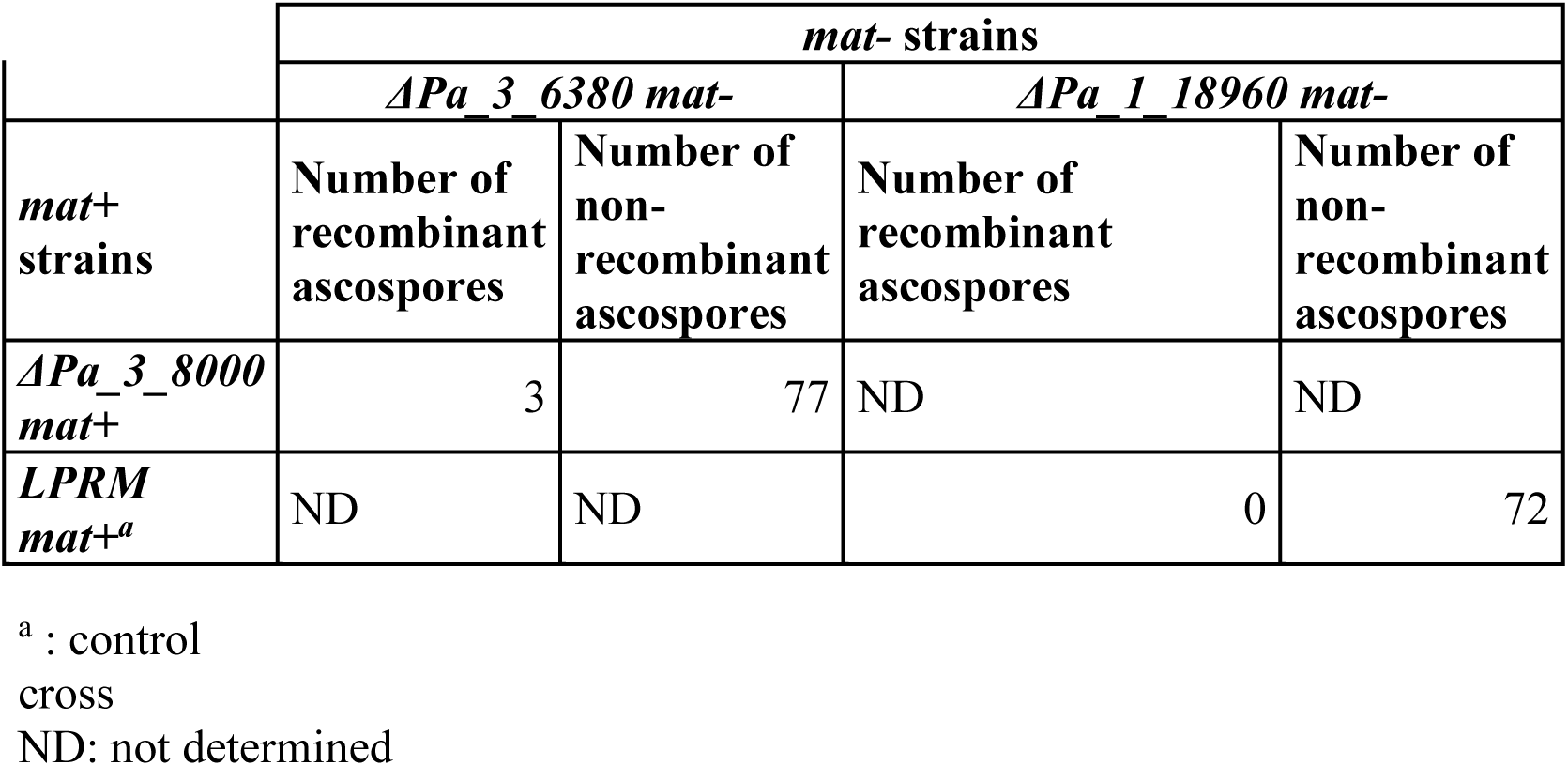
Number of offspring showing recombination between autosomal markers.

As a point of comparison for the expected number of recombination events given the physical distance, we used two other strains, in which the genes *Pa_3_6390* or *Pa_3_8000* have been replaced by a geneticin resistance cassette and a hygromycin resistance cassette, respectively (Grognet, P, unpublished and Debuchy, R, unpublished) (Figure 4B). These two genes are 595 kbp apart on chromosome 3, which is a similar distance to that between the *MAT* locus and *Pa_1_18960* (used to detect recombination in the cross with the *LPRM* strain). By crossing Δ*Pa_3_6390* with Δ*Pa_3_8000,* recombination events between the two genes will yield progeny either resistant to both antibiotics or sensitive to both, allowing detecting recombination events. Among the 80 homokaryotic spores tested, we detected three recombination events (3.75%). A cross of *LPRM* with Δ*Pa_1_18960* was made at the same time and no recombination events between the resistance gene and the *MAT* locus were detected in the 72 homokaryotic spores recovered. Pooling progenies of the three identical crosses involving the *LPRM* strain (*LPRM* x *ΔPa_1_18960 mat-)* indicates that the proportion of recombining offspring in progenies of *LPRM* is significantly different from the proportion in control crosses (significance threshold of 0.05) (Table 3). A similar percentage of recombinants (3.75%) should indeed have led to ca. 11 recombination events among 294 spores between two loci 584 kbp apart on chromosome 1 if recombination had been restored. The results show that the difference in sequences in the *MAT*-proximal region is not responsible for the lack of recombination.

**Table 3:** Statistical comparisons (Fisher’s exact test) using a 2×2 contingency test between proportions of recombinant progenies in crosses involving the *LPRM* strain and a control cross.

| Crosses | Total number of recombinant ascospores | Total number of non-recombinant ascospores | p-value |
| --- | --- | --- | --- |
| **Crosses involving the *LPRM* strain** |  |  |  |
| *LPRM mat+* x *ΔPa_1_18960 mat-* | 1 | 293 | 0.0086 |
| **Control cross** |  |  |  |
| *ΔPa_3_6380* x *ΔPa_3_8000* | 3 | 77 | N.A<sup>a</sup> |

<sup>a</sup>: N.A : not applicable

### What are the molecular mechanisms leading to recombination suppression?

The finding that rendering homozygous the reference strain of *P. anserina* in the *MAT*-proximal region did not restore recombination indicates that sequence divergence alone is not the mechanism responsible for the lack of crossing-overs in this region, nor any *cis*-acting recombination modifiers that would need to be heterozygous to act. It also suggests that sequence divergence is rather a consequence than a cause of recombination suppression.

We replaced the *MAT* locus in a *mat-*genomic background, but we did not perform the reversed experiment by introducing the *mat-*idiomorph in the *mat+* genomic background. Therefore, we cannot exclude that whatever triggers recombination suppression is set up by a *mat*-specific allele that would have a dominant effect in a regular cross (*i.e.* with *mat*- and *mat*+ sequences in the *MAT*-proximal region). However, this hypothesis is very unlikely and there is no obvious candidate sequence for such a role identified in this region (Grognet, Bidard, *et al*. 2014).

The *MAT*-proximal region displays 98.44% of sequence identity between the two mating types in *P. anserina* (Grognet, Bidard, *et al*. 2014; Hartmann, Ament-Velásquez, *et al*. 2021). In crosses between *Podospora* species showing about the same divergence, recombination occurs normally. For example, *P. anserina* and *Podospora comata* display 98% sequence identity genome wide (Boucher *et al*. 2017; Ament-Velásquez *et al*. 2024) but a cross between these two species yields normal recombination rates in the progeny (Espagne *et al*. 2008; Grognet, Lalucque, *et al*. 2014), supporting our conclusion that the sequence divergence observed across the *MAT*-proximal region do not cause recombination suppression.

A single recombination event in the *MAT*-proximal region has been detected in the cross involving the *LPRM* strain. Such a rare event can be expected among a large progeny even in a cross involving strains with wild-type *MAT*-proximal regions (Contamine *et al*. 1996). We have shown that, in a recombination-prone region, the number of recombination events was much higher. In some animal sex chromosomes too, rare events of recombination can occur, as reported for example in frogs (Rodrigues *et al*. 2018). These events have important evolutionary consequences, allowing purging deleterious mutations, regularly “rejuvenating” sex chromosomes (Rodrigues *et al*. 2018).

Because the two mating-type chromosomes are collinear (Grognet, Bidard, *et al*. 2014), inversions or other genomic rearrangements are not responsible either for the recombination cessation. Future studies could investigate epigenetic marks, such as methylation and histone modification, to investigate whether the non-recombining region displays particular patterns that could explain recombination suppression. This would raise the question of what target the chromatin modification specifically to that region.

This study shows the assets of fungi for testing hypotheses on sex-related chromosomes, being experimentally tractable organisms. Of course, the findings on *P. anserina* do not exclude that neutral divergence can cause recombination suppression in other organisms, but it shows that other causes than mere divergence or inversions can suppress recombination. This conclusion is in agreement with previous experiments which demonstrated that even collinear mating-type chromosomes do not recombine in *N*e*urospora tetrasperma* (Jacobson 2005). In *Microbotryum* fungi, young regions without recombination on mating-type chromosomes can also be collinear and with low levels of divergence (Branco *et al*. 2017). Taken together, these experiments emphasize the role of trans-acting inhibitory factors, and tone down the role of chromosome inversions and rearrangements in being the initial proximal cause of recombination suppression, at least in fungi.

The mechanism of recombination suppression and the identification of the *trans*-acting inhibitory factors remain elusive, except in a very few species. Recent progress has been done in *Lachancea klyuvery* and *Chlamydomonas reinhardtii*. In *L. klyuvery*, the absence of recombination in a region of 1 Mb encompassing the mating types was correlated with the absence of synaptonemal complex and meiotic proteins required for recombination (Legrand *et al*. 2024). However, whether this absence relies on the presence of an inhibitory factor or the absence of an activating factor is yet unknown. DNA cytosine methylation was shown to suppress meiotic recombination in the ∼ 300 kbp sex-determining region of the green alga *Chlamydomonas reinhardtii* (Ge *et al*. 2024). The effect of cytosine methylation on the formation of the synaptonemal complex and recruitment of Spo11 has not been investigated. Therefore, communalities for recombination suppression in sex- or mating-type determining regions are still elusive. In *P. anserina*, no cytosine methylation was detected on DNA extracted from mycelium (Bewick *et al*. 2019 Feb 18). However, PaRid, a putative *de novo* DNA-methyltransferase (not related the *C. reinhardtii* DNMT1) is required for reproduction in this species (Grognet *et al*. 2019). The *PaRid* mutant fails to isolate a pair of *mat+* and *mat-*nuclei from the multinucleated cell into a dikaryotic cell to form the future ascogenous hyphae. A seducing hypothesis that arises from this observation is that a transient DNA methylation, reminiscent of the imprinting phenomenon found in animals, would be required first for nuclei identity and later for the regulation of recombination. Interestingly, genetic analyses showed that, in the rare asci where the *MAT* locus undergoes first-division segregation (instead of second-division segregation), the proportion of double crossovers is higher than what is observed in normal asci and some recombination events in the *MAT*-proximal region can occur (Contamine *et al*. 1996). This observation fits with the hypothesis of an epigenetically based recombination regulation. Indeed, the recombination profile is drastically changed and recombination events are frequent when the regulating epigenetic marks are not properly set up.

### Why did recombination suppression evolve?

Regarding the evolutionary causes of recombination suppression around the mating-type locus, sexually antagonistic selection cannot apply to fungi, as male or female functions are not associated to mating types, and there is little trait associated to mating types (Bazzicalupo *et al*. 2019; Hartmann, Ament-Velásquez, *et al*. 2021). This is especially the case in *P. anserina*, in which any role of mating types in determining male and female functions has been discarded (Grognet *et al*., 2014). In *N. tetrasperma*, a previous study proposed that mating-types control feminization and masculinization (Samils *et al*. 2013), but this hypothesis was later discussed (Grognet, Bidard, *et al*. 2014). Furthermore, mating types are expressed at the haploid stage in fungi, and there is virtually no haploid phase in *P. anserina*, during which the two mating types could be selected to behave differently *i.e.*, be subjected to sexually antagonistic selection. A hypothesis that can apply to fungi is the selection of non-recombining fragments that carry fewer deleterious mutations than average in the population, and that are associated to permanently heterozygous loci, which shelter the few deleterious mutations they harbor (Jay *et al*. 2024). A line of evidence supporting this hypothesis is that recombination is suppressed around the mating-type locus only in fungi having an extended dikaryotic (diploid-like) life stage (Jay *et al*. 2024). In Ascomycetes in particular, recombination suppression around the mating-type locus has repeatedly evolved associated with a prolonged dikaryotic stage, which is consistent with an effect of deleterious mutation sheltering (Menkis *et al*. 2008; Vittorelli *et al*. 2023; Jay *et al*. 2024).

## Data availability

The *LPRM* genome sequence is available at http://www.ncbi.nlm.nih.gov/bioproject/1191219 (BioProject ID: PRJNA1191219).

## Acknowledgements

We thank Fanny Hartmann, Daniel Jeffreys, Fabienne Malagnac and Lou Guyot for helpful discussions.

## Author contributions

P.G. and T.G. conceived the experiments. R.D. and P.G. generated the mutant strain. P.G. performed and analyzed crosses, phenotypes and sequences. P.G. and T.G. wrote the paper. T.G. obtained funding.

---

**Figure sup. 1:** Comparison of read coverage at the three loci with putative deletions. Alignment of reads from *LPRM* whole genome sequencing is shown in the lower track. The two upper tracks show the alignment of reads from two controls which should cover the whole genome (namely Input and Mock) of a previous ChIP-seq experiments (Carlier *et al*. 2021). The three putative deleted sequences are devoid of reads in all experiments.

