## Supplementary figures and images for "Genetic differentiation in the *MAT*-proximal region is not sufficient for suppressing recombination in *Podospora anserina*"

### Figure S1

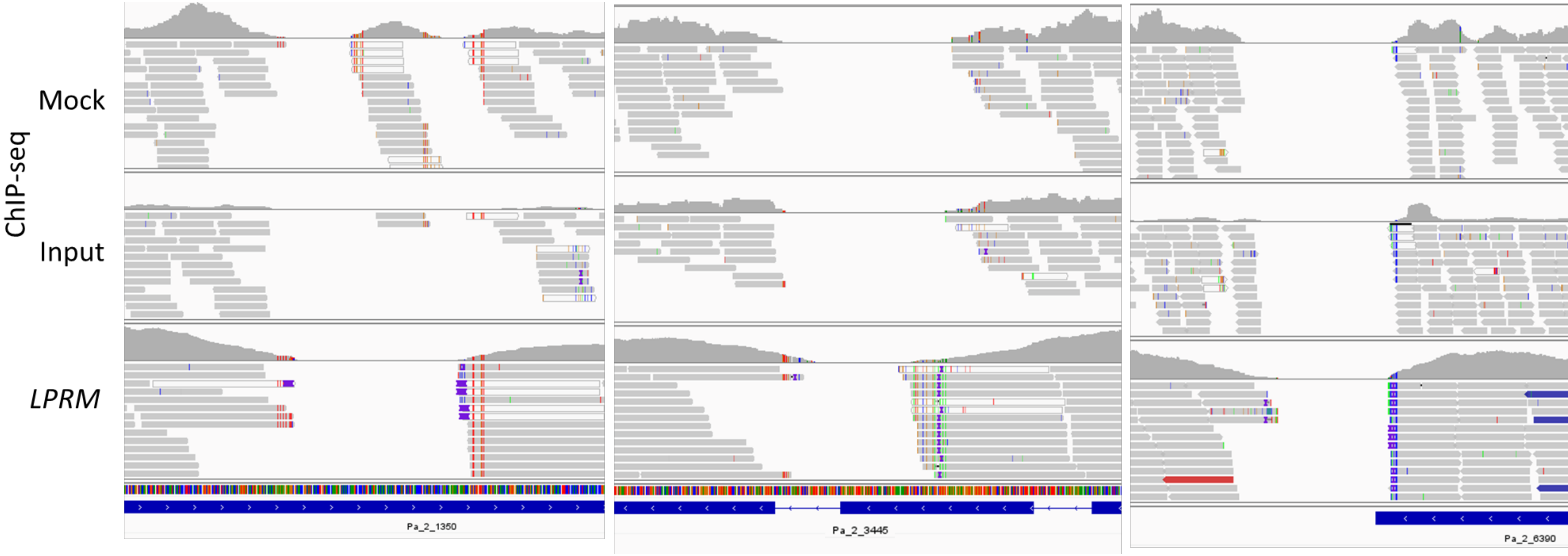
